# A Bayesian Network-Based Framework for Causal Cancer Drug Target Discovery Integrating Patient and Cell Line Data

**DOI:** 10.64898/2026.07.13.736676

**Authors:** Seok Hwan Yoon, Ye Rin Park, Hyun Uk Kim

**Affiliations:** Department of Chemical and Biomolecular Engineering, Korea Advanced Institute of Science and Technology (KAIST), Daejeon 34141, Republic of Korea; Graduate School of Engineering Biology, Department of AX, and BioProcess Engineering Research Center, KAIST, Daejeon 34141, Republic of Korea

## Abstract

Current approaches to cancer drug target discovery face two key limitations: poor translation of cell line-derived targets to patient tumors, and the lack of causal explanation of the regulatory mechanisms underlying target prioritization. Here we present BayesTx (Bayesian Therapeutics target discovery), a Bayesian network framework that integrates patient transcriptomics data with cell line data to identify causal therapeutic targets in cancer. BayesTx projects both data domains into a shared biological space of pathway and transcription factor activities, learns domain-specific causal graphs, and merges them through weighted edge aggregation with bootstrap consensus filtering. Do-simulation on the consensus network quantifies the causal effect of each transcription factor on cancer cell viability. Applied to breast cancer using TCGA-BRCA (The Cancer Genome Atlas breast cancer cohort) and DepMap (Cancer Dependency Map) datasets, the framework ranked 47 transcription factors by predicted causal impact, with gene-level targets further derived through regulon-based propagation. Top-ranked transcription factor (TF) targets were independently supported by survival analysis in external cohort data and pharmacogenomic drug response associations. Overall, BayesTx demonstrates that cross-domain Bayesian network modeling can bridge patient and cell line data to systematically identify causal therapeutic targets in cancer.

## 1. Introduction

Cancer is characterized by extensive molecular heterogeneity, making systematic identification of therapeutic targets from genomic data a persistent challenge^1^. Large-scale resources such as TCGA^2^ (The Cancer Genome Atlas) and DepMap^3^ (Cancer Dependency Map) have profiled thousands of tumors and cell lines, yet characterizing molecular alterations alone is insufficient to identify causal drivers of tumor cell survival. Current approaches to cancer drug target discovery face two fundamental limitations. First, conventional strategies have largely depended on cell line-based screenings^4,5^, but targets identified in vitro frequently fail to translate to patient tumors due to differences in biological context such as tumor microenvironment^6^. Second, existing methods employ correlation-based or black-box models that do not reveal causal pathways^7^. Recent efforts such as TRANSACT^8^ (Tumor Response Assessment by Nonlinear Subspace Alignment of Cell lines and Tumors) and TCGA ^9^ have attempted to bridge the gap between cell line and patient data, but both project one domain into the other’s feature space rather than constructing an integrated causal model encoding directional regulatory relationships.

Bayesian networks (BNs) offer a systematic approach to address these challenges. BNs are directed acyclic graphs (DAGs) that learn causal structure from observational data and support interventional reasoning through do-calculus^10–12^. They have been applied in biological contexts such as gene regulatory network inference^13–15^, yet their use for systematic drug target discovery remains limited. Existing BN merging methods aggregate networks from the same data type to increase statistical power^16^, but are not designed to integrate structurally heterogeneous domains such as patient tumors and cell lines.

Such a cross-domain framework also demands a target class that bridges transcriptional regulation and therapeutic intervention in cancer. Among candidate target classes, transcription factors (TFs) are particularly suited for causal prioritization. TFs function as master regulators driving cancer phenotypes^17,18^, and recent advances in proteolysis targeting chimeras (PROTACs) and molecular glue degraders have shifted the druggability paradigm of TFs^19,20^. TF activity estimates have also been shown to enhance pharmacogenomic response prediction^21^, establishing TFs as both causally important and increasingly druggable targets in oncology.

Here, we present BayesTx (Bayesian Therapeutics target discovery), a generalizable framework for causal drug target discovery. BayesTx projects patient and cell line RNA-seq data into a shared biological space of pathway and TF activities, learns domain-specific graphs via the NOTEARS (Non-combinatorial Optimization via Trace Exponential and Augmented Lagrangian for Structure Learning) algorithm^22^, and merges them into a cross-domain consensus network through weighted edge aggregation with bootstrap consensus filtering. Do-simulation on the consensus network quantifies the causal effect of TF perturbation on cell viability via an average causal effect (ACE) score. As a proof of concept, we apply BayesTx to breast cancer, ranking all 47 TFs by predicted causal impact and highlighting the top 20 candidates. These predictions were independently validated at the TF level through survival analysis and pharmacogenomic correlations, and at the gene level through CRISPR gene dependency data. Overall, our framework provides a systematic and scalable approach for causal target discovery in cancer, bridging observational and interventional datasets to prioritize biologically meaningful and clinically actionable therapeutic candidates.

## 2. Materials and methods

### 2-1. Data preprocessing

CCLE (Cancer Cell Line Encyclopedia) RNA-seq expression data (TPM) was obtained from the DepMap portal (CCLE 2019 release; file: CCLE_RNAseq_rsem_genes_tpm_20180929). Breast cancer cell lines were selected by retaining samples whose lineage annotation contained “BREAST”. Ensembl gene IDs were standardized by removing version suffixes (e.g., ENSG00000000003.10 to ENSG00000000003). Missing expression values were imputed as zero, and all values were cast to 32-bit floating point. Expression values were log₂-transformed using log₂(TPM + 1) to stabilize variance and align with the TCGA preprocessing pipeline. The resulting expression matrix comprised 57820 genes across 51 breast cancer cell lines in log₂-TPM units.

CRISPR-Cas9 gene dependency scores (Chronos-corrected gene effect) were obtained from the DepMap portal (DepMap Public 25Q3; file: CRISPRGeneEffect.csv). Cell line metadata (DepMap Public 25Q3; file: Model.csv) was used to filter breast-lineage models and map DepMap model identifiers to standardized cell line names matching the CCLE RNA-seq column format. Genes with missing dependency values were removed, yielding a final matrix of 17,204 genes across 53 breast cancer cell lines.

Patient-level gene expression data was obtained from TCGA-BRCA (The Cancer Genome Atlas breast cancer cohort) via the GDC (Genomic Data Commons) portal. From individual RNA-seq quantification files (augmented STAR gene counts), unstranded TPM values were extracted. Ensembl gene IDs were standardized by removing version suffixes, and non-gene summary entries were excluded. For duplicate gene IDs after version removal, TPM values were summed to obtain unique gene-level expression. Individual samples were concatenated into a unified 16,391 genes across 1,111 samples, with missing values imputed as zero, followed by log₂(TPM + 1) transformation.

### 2-2. Network node design and calculation

The causal graph architecture comprises the following node types: pathway activity nodes (layer 1), transcription factor activity nodes (layer 2), and domain-specific outcome nodes (clinical node and viability node). The layers correspond to successive levels of biological regulation, capturing the information flow from signaling pathways through transcriptional regulation to phenotypic outcomes.

#### Gene identifier conversion

To enable pathway and TF activity scoring from Ensembl-indexed expression data, gene identifiers were converted from ENSG IDs to HGNC symbols using the MyGene.info API. The conversion prioritized human protein-coding genes and selected the highest-confidence match per ENSG ID. Where multiple ENSG IDs mapped to the same symbol, expression values were averaged. This yielded symbol-indexed expression matrices compatible with pathway and regulon gene set definitions.

#### Layer 1: Pathway activity scoring

Two methods were employed to capture signaling pathway activity from gene expression:

PROGENy^23^ (Pathway RespOnsive GENes) activity scores were computed using the decoupleR^24^ package. For each of 14 cancer-related pathways (Androgen, Estrogen, EGFR, Hypoxia, JAK-STAT, MAPK, NFkB, p53, PI3K, TGFb, TNFa, Trail, VEGF, WNT), the top 500 genes ranked by absolute pathway-association weight were selected from the PROGENy model. Pathway activity was estimated using a weighted mean, computed as Σ(w·x) / Σ|w|, where w denotes the pathway-gene association weight and x denotes log₂-transformed expression values. A multivariate linear model (MLM) is also available as an alternative scoring method. Scores were z-score normalized per pathway across samples.

Hallmark gene sets from the MSigDB^25^ (Molecular Signatures Database v2023.2) were scored using ssGSEA (single-sample Gene Set Enrichment Analysis) via GSEAPy with rank-based sample normalization. Enrichment scores were computed for 50 hallmark pathways per sample, capturing functional pathway states independent of predefined gene weights. Gene sets were filtered to retain pathways containing 10 to 500 genes, and resulting scores were z-score normalized per pathway across samples.

#### Layer 2: Transcription factor activity scoring

Transcription factor regulatory activity was inferred from regulon expression using two curated regulatory networks:

DoRothEA^21,26^ (Discriminant Regulon Expression Analysis) regulons were obtained via the decoupler package^24^, filtered to confidence level A (highest-confidence interactions), and required a minimum of 25 target genes per TF. The framework supports multiple TF activity estimation methods - ULM (Univariate Linear Model), VIPER (Virtual Inference of Protein-activity by Enriched Regulon analysis), WMEAN (weighted mean), and GSVA (Gene Set Variation Analysis) - selectable via command-line parameter. ULM estimates TF activity as the t-statistic from regressing regulon weights against expression; VIPER computes a weighted enrichment score over the ranked expression signature; WMEAN calculates the weight–expression dot product normalized by the sum of absolute weights (Σwᵢxᵢ / Σ|wᵢ|); and GSVA separately scores activating and repressing target gene sets per sample, subtracting the latter from the former to yield a signed enrichment. Scores were z-score normalized per TF across samples.

CollecTRI^27^ (Collection of Transcriptional Regulatory Interactions) regulons were obtained via the decoupler package. When the CollecTRI source annotation file was provided, interactions were filtered for curated evidence, requiring support from at least one of seven literature-curated databases (TRRUST, HTRI, TFactS, SIGNOR, NTNU Curated, Pavlidis2021, and DoRothEA_A). TFs with fewer than 25 target genes were excluded. TF activity scoring methods were identical to those described for DoRothEA (ULM, VIPER, WMEAN, GSVA).

#### Clinical nodes (patient domain)

Clinical covariates were obtained via the GDC REST API by querying TCGA-BRCA RNA-seq files (STAR-Counts workflow, TSV format). For each file, the sample-level UUID was extracted from the file name using regular expression matching, and demographic variables (age, gender, and race) were retrieved from the associated case metadata. Duplicate entries were removed by retaining the first occurrence per UUID. The resulting clinical matrix was linked to the expression data using file-level UUIDs as shared identifiers.

For Bayesian network construction, clinical variables were classified as continuous or categorical based on data type: age was treated as a continuous variable, while gender and race were encoded as integer codes (categorical variables with ≤10 unique levels). One sample with a missing age value was excluded, reducing the cohort from 1,111 to 1,110 patients. These clinical variables were concatenated with PROGENy pathway activity scores (14 pathways) and DoRothEA TF activity scores (47 TFs) across the intersection of matched samples, yielding a unified input matrix of 1,110 patients with 64 variables. All variables were standardized using z-score normalization prior to structure learning.

#### Viability node (cell line domain)

A global cell viability score was derived from CRISPR-Cas9 gene effect data to serve as the outcome node in the cell line domain-specific graph. Genes and cell lines with more than 30% missing values were excluded, and remaining missing values were imputed with gene-wise medians. Genes with zero variance were removed. The first principal component (PC1) was extracted from the gene-wise z-scored gene effect matrix and oriented so that higher values correspond to greater cell viability. The resulting score was z-score normalized across cell lines. This viability score was merged with PROGENy and DoRothEA activity scores across the 34 cell lines common to all three matrices, producing an input matrix of 62 variables, z-score normalized prior to structure learning.

### 2-3. Domain-specific causal graph learning

Using the calculated activity scores of both pathway and TF from 1,110 patients and 34 cell lines, we constructed separate causal graphs for the patient and cell line domains using NOTEARS algorithm^22^. NOTEARS uses continuous optimization to search for the optimal graph structure, which reduces the operation time needed to build every separate candidate structure and score them, as used in conventional score-based learning^28^.

#### Node composition and directional constraints

Each domain graph is composed of three kinds of nodes as noted in section 2-2. For the patient domain, clinical nodes (domain-specific layer), pathway activity nodes (layer 1) and TF activity nodes (layer 2) were used, whereas for the cell line domain, pathway activity nodes (layer 1), TF activity nodes (layer 2) and viability nodes (domain-specific layer) were used. Two domains were designed to share the same pathway activity and TF activity nodes, which are used to merge the graph afterwards (section 2-4).

To encode prior biological knowledge into the structure learning, a blacklist mechanism was applied post-hoc to the NOTEARS weight matrix. Weights of the edges in the blacklist were set to zero regardless of the data-driven solution, prohibiting biologically implausible connections. The blacklist constraints are of two types: automated layer-direction constraints, derived from the assumed hierarchical relations, and user-defined blacklist edges which can be optionally used to further restrict the edge connection based on prior biological knowledge. For the automated layer-direction constraints, Pathway to Clinical edges in the patient domain and outgoing edges from the viability node were prohibited, reflecting viability node’s role as the terminal outcome node (S1). Also, as a baseline model of this study, TFs are placed downstream of pathway activity nodes, reflecting their role as executors of pathway-level transcriptional programs (TF to Pathway edges prohibited)^23,26^. However, to consider the biological relationship of TFs affecting the pathway-level activity^29^, graph structures without directional constraints were additionally tested (section 2-5). A whitelist option is also available in the implementation. However, as it operates as a post-hoc weight moderation to non-existent edges rather than a structural constraint enforced during optimization, its effect in the domain-specific graph structure is limited.

#### Edge weight threshold

Following the post-hoc prior knowledge application, edges were retained in the domain-specific graph based on an absolute weight threshold. In detail, edges were preserved in the thresholded graph provided their absolute weight satisfied the condition |*w_ij_*| ≥ τ, where *w_ij_* represents the NOTEARS-optimized coefficient connecting node i to node j. The user-specified parameter τ (defaulting to 0.001) served as the filter for structural importance. Crucially, the non-thresholded continuous matrix served as the primary input for subsequent cross-domain graph integration, ensuring the preservation of subtle structural dependencies before final graph pruning.

### 2-4. Cross-domain graph merging

Domain-specific causal graphs from each patient domain and cell line domain share the pathway activity nodes and the TF activity nodes in common. Using these nodes, the two graphs can be linked together. Since each domain has a different sample size and data distribution, weight from each domain went through max-abs scaling. Afterwards, various merging methods were tested and compared.

For the edges coexisting on both domain graphs with same weight signs, the weights in the merged graph were calculated using weighted mean. The weighted mean uses the merge ratio (α) of each domain while calculating the final merged weight. This merge ratio of two domains can be modified by the user, with two numerical values summed to 1 (α_1_ + α_2_= 1). in the equation is the scaled weight of domain k. For edges with conflicted weight signs, (i.e., edge A to B in patient domain and B to A in cell line domain), conflict was resolved by removing edges with conflicting signs. The method explained was selected after the comparison of three different merging methodologies. The resulting graph was further constrained to be acyclic by iteratively removing the weakest edge within any remaining cycle. On testing the merge method, a merge ratio sweep was performed (α_1_ : α_2_ = 1:9 ∼ 9:1) to get the best merge ratio with highest Cross-validated Log-likelihood (CVLL). Mean Square Error (MSE) was also calculated for each run. CVLL and MSE were calculated based on the Gaussian structural equation model (SEM), with 5-fold cross validation.

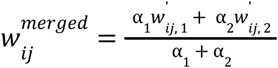

### 2-5. Selection of baseline model

Sixteen module configurations were evaluated by combining pathway activity sources (PROGENy, Hallmark, Hallmark filtered to ≥90% gene coverage), TF regulon sources (DoRothEA, CollecTRI), TF activity estimation methods (weighted mean, ULM, VIPER, GSVA), and layer constraint modes (closed or opened). Each configuration was assessed using four cross-validated Gaussian SEM metrics summed across both domains: CVLL per sample, CVLL per observation, MSE per sample, and MSE per observation. Per observation metrics normalize per sample metrics by both sample and node count, enabling fair comparison across configurations with different numbers of variables.

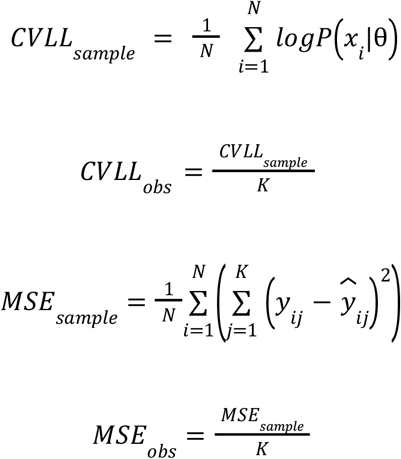

where N is the number of samples, K is the number of nodes (variables) in the model, *x_i_* is the observed data vector for sample i, θ is the set of SEM parameters, *y_ij_* and *ŷ_ij_* are the observed and predicted values for sample i at node j, respectively. Domain-specific scores were summed across patient domain (TCGA-BRCA) (N = 1,110, K = 64) and cell line domain (DepMap) (N = 34, K = 62) to yield a single comparative metric per configuration.

Among the sixteen configurations compared, the PROGENy-DoRothEA-weighted mean configuration with the baseline constraint (closed) achieved the highest CVLL per sample (−128.360), indicating the most consistent explanatory power across both domains. The same pathway-TF scoring method combination under the optional constraint (opened) showed the lowest MSE per sample (54.327), while the Hallmark-DoRothEA-weighted mean configuration under the optional constraint (opened) ranked first in both log-likelihood per observation (−1.899) and MSE per observation (0.802). The closed-layer constraint places TFs downstream of pathways, consistent with their biological role as executors of pathway-level transcriptional programs. Thus, the PROGENy-DoRothEA-weighted mean configuration was selected as the baseline configuration for all subsequent analyses including do-simulation, target prioritization, and validation.

### 2-6. Bootstrap stability and consensus network

To ensure stability of the merged graph, the whole process of obtaining the merged graph, from domain-specific causal graph learning to cross-domain graph merging, was repeated. NOTEARS-based structure learning produces a fixed solution for a given dataset. However, the learned graph structure can vary with the changes in sample composition, especially in DepMap since the number of breast cancer cell lines is limited. To obtain a structurally robust graph, we applied a bootstrap (B = 200) procedure across the full pipeline. In each of the bootstrap replicates, samples were gathered with replacement independently from patient domain (TCGA-BRCA) and cell line domain (DepMap), domain-specific BN structures were learned, and merged. For every single edge that had occurred at least once throughout the bootstrap process, bootstrap probability was calculated. This probability is the probability of each edge appearing in the graph structure, which can be interpreted as the stability of an edge in the given dataset.

To obtain the final bootstrap consensus network structure, the structurally robust merged graph retained for downstream causal inference, we applied a threshold to the bootstrap probability of edges. Only edges with bootstrap probability over 0.5, which means that the edges were present in over 100 of 200 bootstrap replicates, were retained. Remaining edges went through DAG constraints to ensure acyclicity.

### 2-7. Target ranking via do-simulation

#### Target ranking via do-simulation and gene-level extension

From the resulting consensus network, target nodes were ranked by the change of cell viability (Δviability) after a negative intervention (-1σ). To do so, Gaussian SEM was fitted on the final graph structure using node-wise OLS regression on cell line domain (DepMap) data. The average causal effect (ACE) score of each node on viability was estimated via do-calculus intervention, *do*(*Z* = µ*Z* − σ*Z*). We refer to this do-calculus-based intervention procedure as do-simulation.

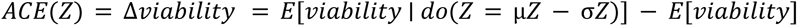

Node rankings were listed by the computed ACE score, with the sign indicating the direction of therapeutic strategy. Nodes with negative ACE can be interpreted as inhibition targets, and nodes with positive ACE can be interpreted as overexpression targets.

As TF and pathway nodes operate at the clustered activity level rather than the gene level, gene-level causal scores were derived by propagating TF-level ACE estimates through the DoRothEA regulon using a confidence-weighted mean, as described in Section 2-2.

#### Mechanistic validation of gene-level causal predictions

To evaluate whether the inferred causal directions align with functional viability patterns, breast cancer cell lines were stratified into high- and low-expression subgroups (top and bottom 30%) for each target gene. Statistical significance of viability differences between groups was determined via Mann-Whitney U test, and overall associations were quantified using Spearman correlation with FDR-adjusted p-values. For each gene, the theoretical viability impact was defined as the product of the BN-derived edge sign and the regulon-specific weight sign. We subsequently assessed sign consistency to verify the degree to which predicted regulatory effects match observed phenotypic outcomes across the cell line domain.

### 2-8. Validation and evidence profiling of target candidates

#### Literature based profiling of target candidates

Conventional literature mining tools in the field of cancer (CancerMine^30^, Open Targets^31^) focus on individual genes. However, since our framework suggests TF-level cancer targets, the literature mining in our framework was designed to assess the biological plausibility of each TF candidate as a guidance. Specifically, target nodes were categorized by the sign of their causal effect on viability: nodes with a negative ACE, where decreased activity leads to viability decrease were treated as oncogene-like candidates and evaluated for inhibition evidence (inhibition targets), while nodes with a positive ACE where decrease activity leads to viability increase were treated as tumor suppressor-like candidates and evaluated for loss-of-function and synthetic lethality evidence (overexpression targets).

For each target, PubMed was queried across three categories: functional evidence, clinical relevance, and drug availability, using search keywords constructed from the target node name and the cancer type (“breast cancer” as default), supplemented with MeSH terms. Literature mining was computed for top-ranked candidates as a supporting annotation, providing an independent assessment of existing evidence for each target rather than serving as a selection criterion.

#### METABRIC survival analysis

To evaluate whether the top-ranked TF candidates identified by the pipeline are associated with clinical outcomes in an independent patient cohort, survival analysis was performed on the METABRIC (Molecular Taxonomy of Breast Cancer International Consortium) dataset^32^. TF activity scores for METABRIC patients were computed from microarray gene expression data using the same DoRothEA-based weighted mean scoring applied in patient domain (TCGA-BRCA), with confidence level A regulon only. Patients were stratified into high- (top 30%) and low-activity (bottom 30%) groups based on each TF’s activity score, and Kaplan-Meier survival curves were generated for overall survival (OS) and relapse-free survival (RFS). Group differences were evaluated using the log-rank test. In addition, Cox proportional hazards regression was performed with TF activity as a continuous covariate to estimate the hazard ratio (HR) and 95% confidence interval, providing a complementary measure of effect size independent of the stratification threshold.

#### GDSC2 drug response association

To assess the therapeutic relevance of top-ranked TF candidates, TF activity scores derived from CCLE breast cancer cell lines were correlated with drug IC50 values from the GDSC2 dataset^33–35^. Cell lines were stratified into high- (top 30%) and low-activity (bottom 30%) groups for each TF, and group differences in ln-transformed IC50 values were assessed using the Mann-Whitney U test with rank-biserial correlation as the effect size measure. FDR correction (Benjamini-Hochberg) was applied independently within each TF across all tested drugs. Associations with FDR < 0.05 were considered significant, and directional consistency with BN-predicted intervention strategy was evaluated: for inhibition targets (positive ACE), higher TF activity was expected to associate with increased IC50, reflecting reduced drug sensitivity; for overexpression or synthetic lethality targets (negative ACE), higher TF activity was expected to associate with decreased IC50.

## 3. Results

### 3-1. Cross-domain Bayesian network integrates patient and cell line RNA-seq data

We developed BayesTx, a cross-domain causal Bayesian network framework that integrates patient-level clinical and transcriptomic data with cell line-level transcriptomic and viability data to identify and prioritize candidate therapeutic targets at the pathway, transcription factor, and gene levels (Sections 2-1–2-7). The BayesTx framework was applied to breast cancer using patient domain (TCGA-BRCA) transcriptomics and clinical data with cell line domain (DepMap/CCLE) transcriptomics and CRISPR dependency data. Structure learning yielded a patient-domain graph of 64 nodes and 521 edges (4,032 before threshold filtering), and a cell line-domain graph of 62 nodes and 497 edges (3,782 before threshold filtering). The smaller scale of the cell line-domain graph reflects the limited number of breast cancer cell lines available in DepMap. To obtain a structurally robust merged network, we applied bootstrap resampling (B = 200), and 2,806 distinct edges were observed at least once. Retaining only edges with bootstrap probability ≥ 0.5 yielded the final consensus network of 64 nodes and 275 edges (Fig. 2h).

**Figure 1.**
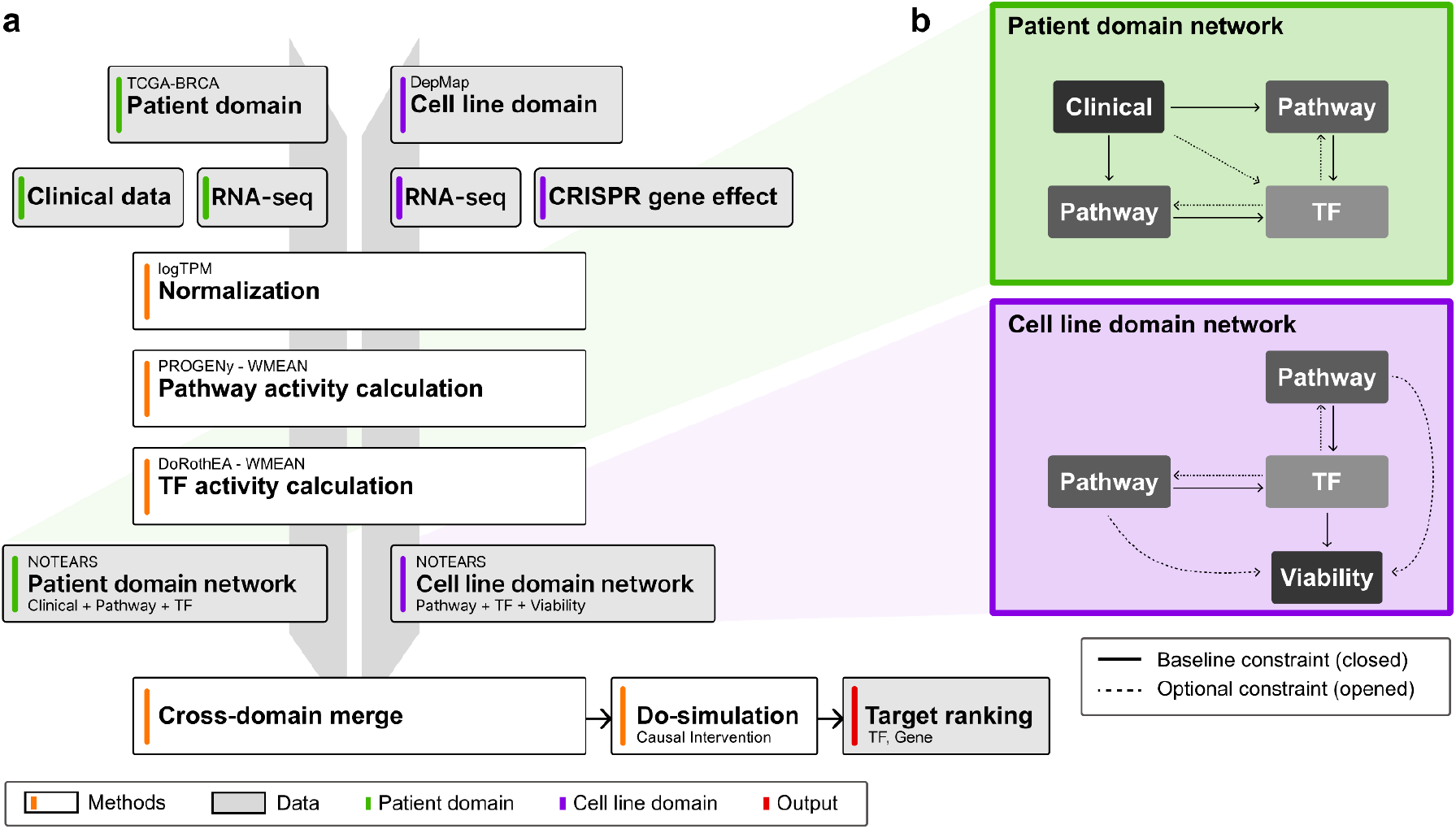
Overall pipeline of BayesTx applied on BRCA and scheme of the output network. **a** The BayesTx framework integrates two domains: a patient domain (TCGA-BRCA clinical and RNA-seq data) and a cell line domain (DepMap CRISPR gene-effect and RNA-seq data). RNA-seq from each domain is independently processed into pathway activity (PROGENy) using weighted mean (WMEAN) and transcription factor (TF) activity (DoRothEA) scores. A domain-specific network is learned for each domain using NOTEARS, and the two networks are merged by retaining edges with concordant signs. Causal effects on cell viability are then quantified by fitting a Gaussian SEM to the merged structure and applying do-simulation; nodes are ranked by the resulting viability change. **b** Structural scheme of the two domain-specific networks: patient domain and cell line domain. Solid arrows indicate the baseline constraint (closed) that restricts directionality to a ‘clinical - pathways - TFs - viability’ cascade, assuming that TF is a downstream regulator of pathways. Dashed arrows indicate the optional constraint (opened) that allows the linkage from TF to pathways, and clinical to TFs, which supports the complicated nature of human regulatory networks.

**Figure 2.**
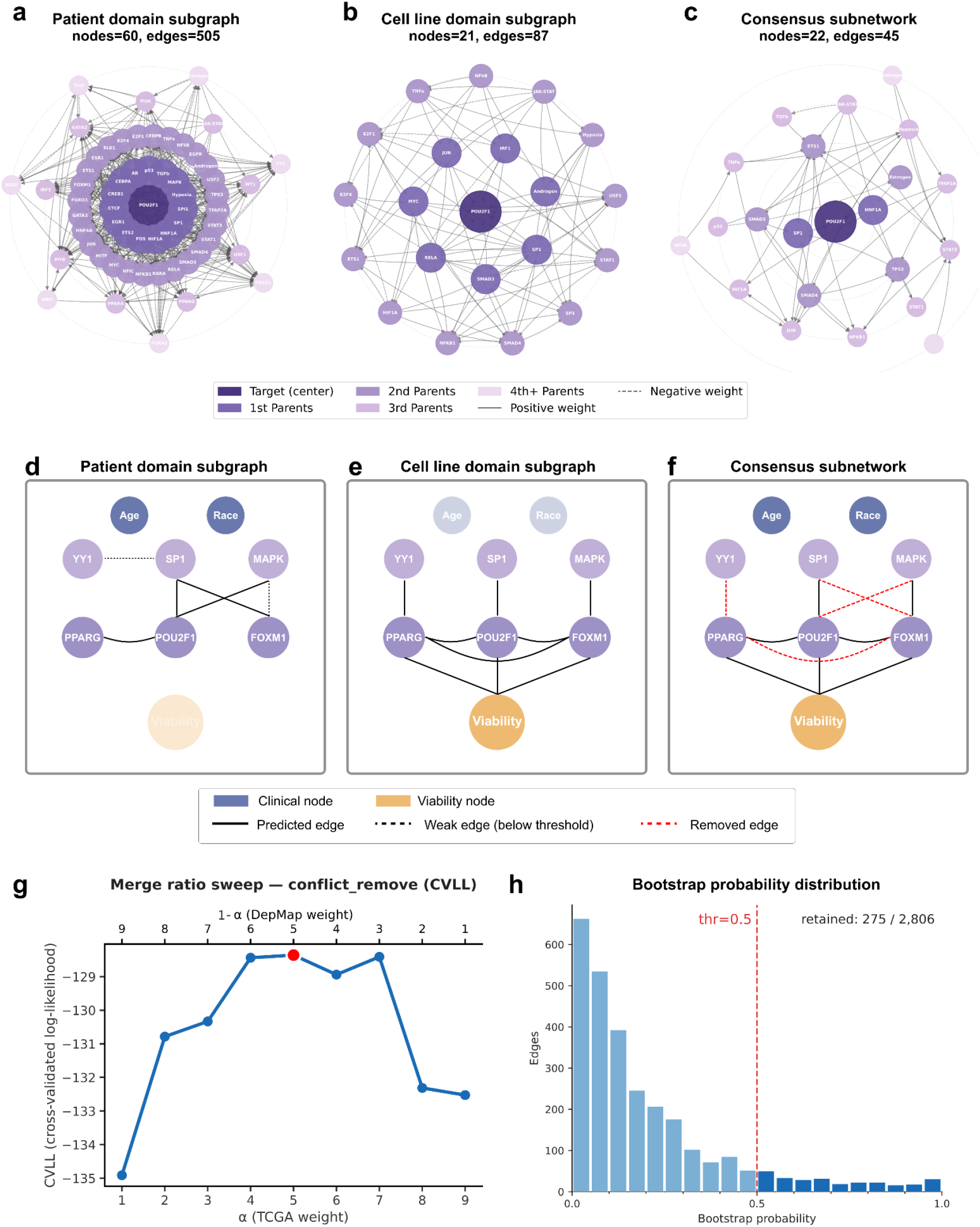
BayesTx pipeline applied to breast cancer (TCGA-BRCA, DepMap): domain-specific Bayesian networks, cross-domain merge and parameter selection. **a-c** Circular subgraphs centered on the POU2F1 TF node. Each panel displays a subnetwork within a fixed depth from the centered node POU2F1, with nodes arranged in concentric rings by hop distance. Node color indicates the layer depth: centered node (dark purple), 1st parents (medium purple), 2nd parents, 3rd parents, and 4th parents (progressively lighter purple). Edge color indicates directionality of regulatory influence: solid black lines represent positive-weight edges and red dashed lines represent negative-weight edges. **a** Patient-domain subgraph learned from TCGA-BRCA RNA-seq and patient clinical data (60 nodes, 505 edges). **b** Cell line-domain subgraph learned from DepMap/CCLE RNA-seq and CRISPR gene-effect data (21 nodes, 87 edges). **c** Consensus subnetwork after cross-domain weighted integration (22 nodes, 45 edges). **d-f** The effect of network merging visualized. Network structure between the same nodes shows different connections within patient domain (**d**) and cell line domain (**e**). Through the merging process, only edges with the same sign on both domains are retained (**f**). **g** Selection of the optimal domain merging ratio via parameter sweep. Cross-validated log-likelihood (CVLL) per sample across a range of merging ratios (α), where α represents the relative weight assigned to the patient domain (TCGA-BRCA). The red dot indicates the optimal α value corresponding to the highest CVLL. The merging ratio at the optimal α was used for all subsequent analyses. **h** Bootstrap probability distribution of merged graph edges. Edges with probability lower than 0.5 were removed from the final consensus network.

To characterize how cross-domain integration shapes the resulting consensus network topology, we compared the domain-specific graphs and the merged consensus network at the global structural level. The merging process substantially reduced network density relative to either individual domain, reflecting the stringency of the sign-concordance filtering applied during integration: an edge survives merging only if it is independently supported by both the patient and cell line domains. This effect can be illustrated by examining the local neighborhood of a representative hub node. The transcription factor POU2F1-centered subgraph is visualized for each domain (Fig. 2a–b) and the consensus network (Fig. 2c) as an example; in the patient domain, POU2F1 was embedded within a structurally rich neighborhood spanning up to four layers, whereas the cell line domain showed a more compact subgraph around the same node, and the consensus network showed the sparsest structure of the three. Comparison of the same node pairs across the two domains revealed partially divergent edge structures (Fig. 2d–e), confirming that patient-level transcriptomic regulation and cell line-level viability-linked regulation capture complementary aspects of breast cancer biology. Following cross-domain integration with the conflict_remove strategy, only edges with concordant sign across both domains were retained, yielding a merged consensus network (Fig. 2f). Also, it is shown that edges in the merged structure have not originated from a single domain.

The merge ratio sweep identified α = 0.5 as the optimal domain weighting (Fig. 2g), indicating that equal contribution from both domains maximized generalization as measured by CVLL. The maximization of the metric at equal weighting suggests that patient domain and cell line domain networks carry complementary, non-redundant information, and that neither domain alone is sufficient to capture the regulatory relationships relevant to breast cancer viability.

## 3-2. Hierarchical do-simulation identifies layer-specific therapeutic targets

### TF-level target identification via do-simulation

To prioritize therapeutic targets at the regulatory level, we applied do-simulation to each pathway and TF node in the consensus network, quantifying the causal effect of inhibiting each node on cancer cell viability (ACE). At the TF level, all 47 TFs in the consensus network were evaluated. Of these, 14 TFs showed negative ACE score (inhibition candidates), 32 TFs showed positive ACE score (overexpression candidates), and 1 TF (CEBPA) showed near-zero effect (Figure 3a). Together, these results show a ranked set of TF-level targets based on the causal structure of the consensus network.

**Figure 3.**
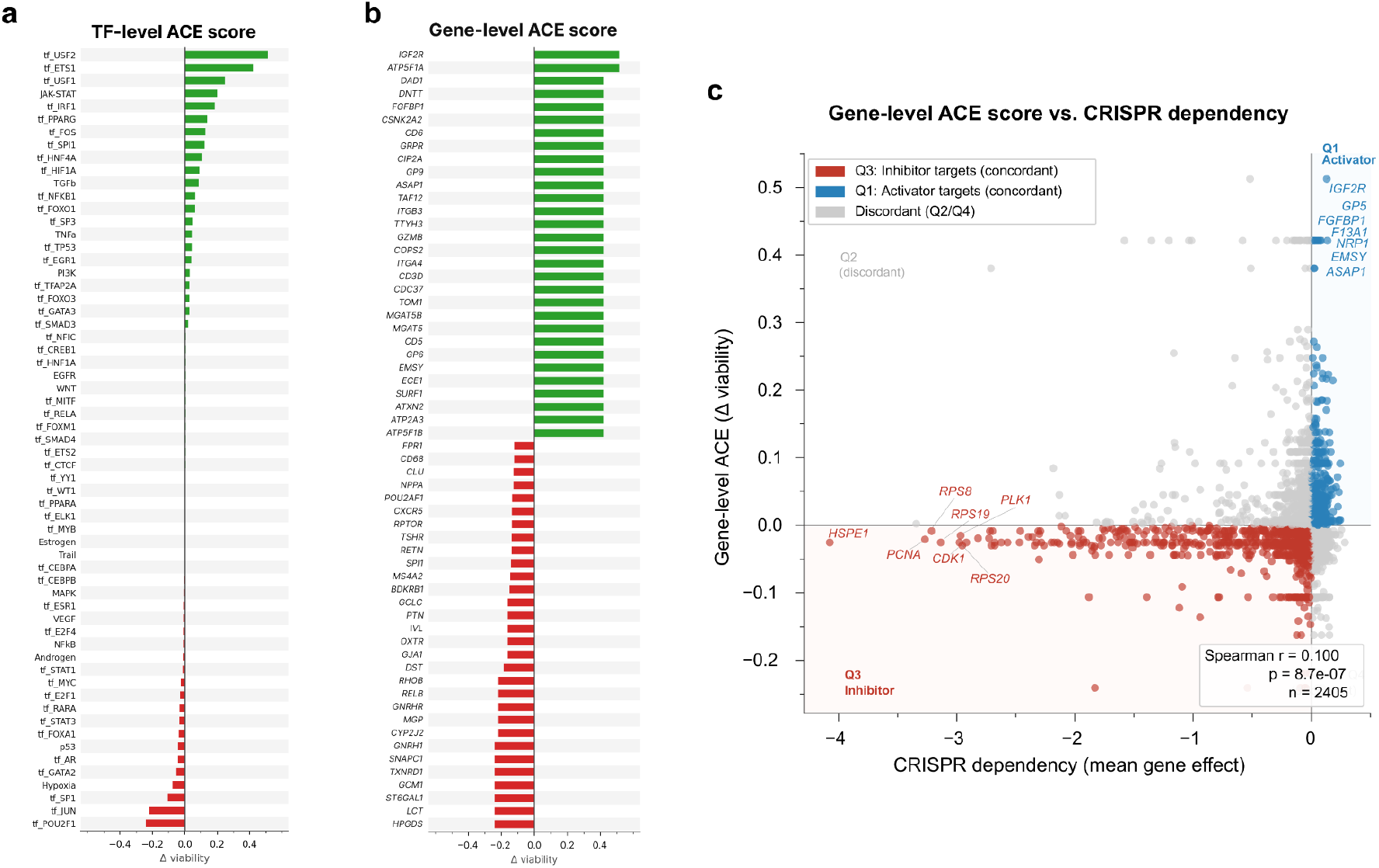
TF- and gene-level target ranked via do-simulation. **a** Pathway- and TF-level ACE scores (Δviability) were estimated by do-simulation (-1**σ**) on the consensus network. Positive values indicate increased viability upon -1**σ** knockdown intervention, and negative values indicate decreased viability upon same intervention. **b** Gene-level ACE scores were computed by propagating TF-level ACE score through the DoRothEA regulon, using signed regulatory connections and confidence levels. **c** Comparison of gene-level ACE scores and CRISPR gene dependency across breast cancer cell lines (n = 53). Gene-level ACE score was derived from TF-level scores and DoRothEA regulon weights. CRISPR dependency values represent the mean gene effect across breast cancer cell lines. Concordant genes in Q1 (ACE-positive, CRISPR-positive) and Q3 (ACE-negative, CRISPR-negative) represent activator target and inhibitor target candidates, respectively.

### Gene-level target scoring via TF regulon-based propagation

To translate TF-level causal scores into gene-level target candidates, TF-level ACE were projected onto regulon genes through DoRothEA using signed regulon weights. For each 2,405 regulon gene, the calculated score reflects the predicted viability impact mediated through its upstream TF (Figure 3b).

### Gene-level predictions are supported by CRISPR gene dependency

Gene-level ACE scores were compared to CRISPR gene effect data of breast cancer cell lines to evaluate whether causal predictions align with functional gene essentiality. Genes in Q3 (negative do-simulation and negative CRISPR gene effect) represent inhibition candidates supported by both causal modeling and experimental essentiality, and genes in Q1 (both positive) represent activator candidates. A total of 924 and 423 genes fell in Q3 and Q1, respectively, out of 2,405 evaluated genes (Figure 3c). Notable Q1 and Q3 genes included core essential genes and progression driving genes. Across all genes, ACE scores showed a significant positive correlation with CRISPR dependency (Spearman ρ = 0.100, *p* = 8.7 × 10⁻⁷, n = 2,405), indicating consistency between BN-derived causal predictions and experimental gene essentiality.

#### 3-3. Computational validation of known and unknown targets

To test the biological relevance of the suggested top-ranked TF candidates, we performed three complementary analyses: survival association in an independent clinical cohort, drug response correlation in an independent cell line dataset, and literature-based evidence quantification.

### Survival analysis

Kaplan-Meier survival analysis was performed on the METABRIC cohort, classifying patients by high and low TF activity scores (top and bottom 30%, Section 2-7). The majority of the top BN-predicted inhibition targets showed consistent prognostic signals across both endpoints, OS and RFS, also in the direction predicted by the BN (Figure 4a).

**Figure 4.**
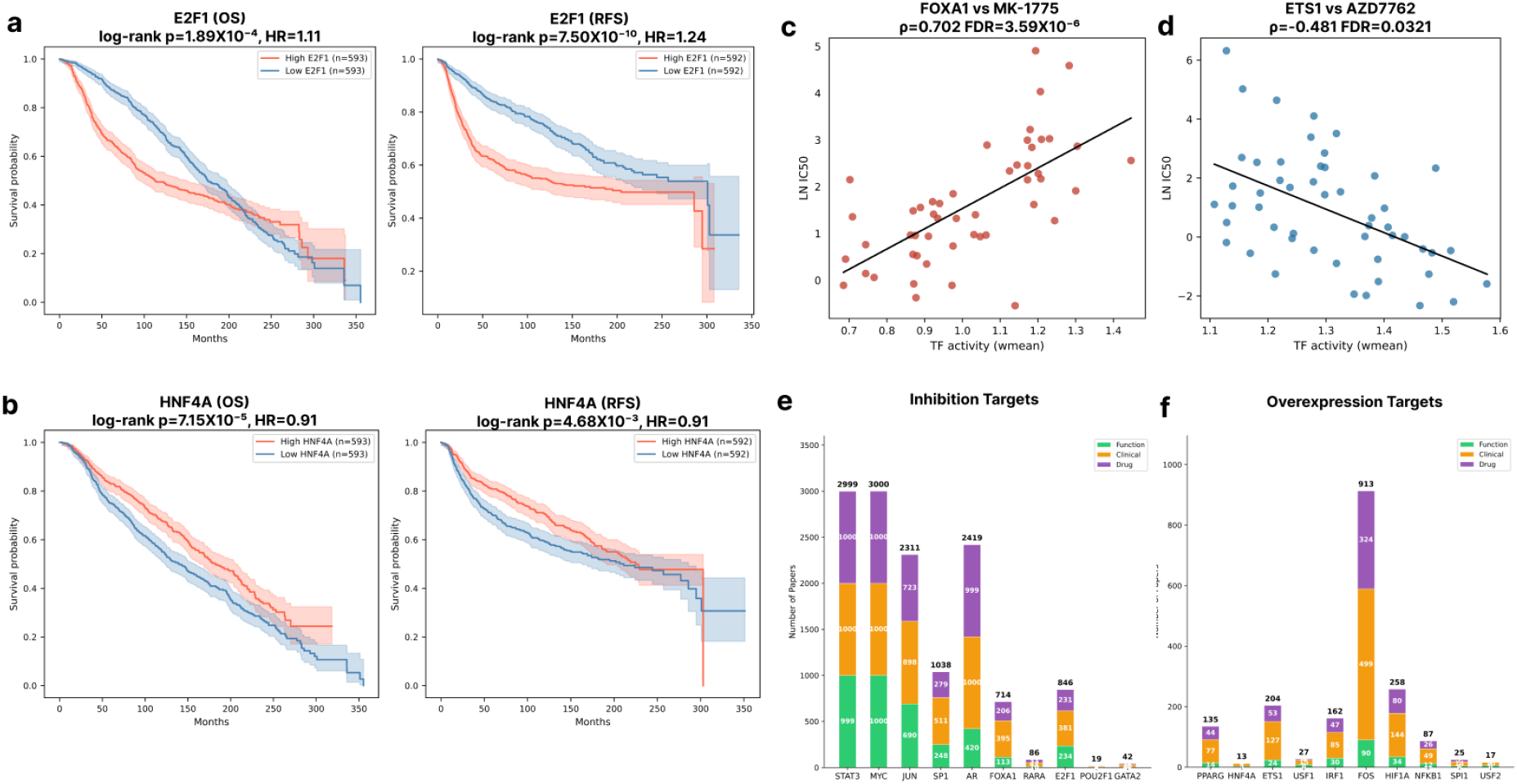
Validation of top ranked TFs. **a** Kaplan-Meier survival curves for E2F1, stratified by high (red) and low (blue) TF activity, for overall survival (OS) and relapse-free survival (RFS). **b** Kaplan-Meier survival curves for HNF4A, stratified by high (red) and low (blue) TF activity, for OS and RFS. Shaded areas represent 95% confidence intervals. P-values were calculated using the log-rank test. **c-d** Scatter plots showing representative TF–drug pairs selected as the top FDR-significant associations with direction consistent with BN prediction within each group: inhibition targets (**c**) and overexpression targets (**d**). Each dot represents a breast cancer cell line. Regression lines are shown with 95% confidence intervals. **c** FOXA1 vs. MK-1775 (ρ = +0.702, FDR = 3.59 × 10⁻⁶) demonstrate that higher TF activity associates with increased IC50, consistent with the BN-predicted inhibition direction. **d** ETS1 vs. AZD7762 (ρ = −0.481, FDR = 0.0321) demonstrate that higher TF activity associates with decreased IC50 for the Chk1/2 inhibitor AZD7762, consistent with the BN-predicted overexpression direction. IC50 values are shown on a natural log scale. **e-f** Stacked bar charts showing the number of PubMed publications retrieved for each TF across three categories: Function (green), Clinical (orange), and Drug (purple), for inhibition targets (**e**) and overexpression targets (**f**). For each TF, literature searches were performed using category-specific query terms. Counts exceeding 1,000 were capped at 1,000 per category (maximum total = 3,000). Numbers above each bar indicate total publication counts.

Among the remaining inhibition targets, several showed statistically significant associations with survival in the direction opposite to BN predictions. These target TFs were each associated with improved rather than worsened survival in groups with higher TF activity (all HR < 1). For target TFs with non-coherent results between drug response and survival analysis, the discrepancy may reflect the difference between cell viability, the outcome modeled by the BN, and overall clinical survival.

For these targets, overall clinical survival integrates additional factors, such as treatment response and tumor microenvironment that are not captured in the do-simulation framework.

Among the top-ranked overexpression targets, HNF4A showed significant associations with both overall survival and relapse-free survival, with higher HNF4A activity associated with improved patient outcome (Figure 4b). This is consistent with the BN-predicted direction, in which higher HNF4A activity decreases cancer cell viability and, correspondingly, was associated with better clinical outcome, supporting its classification as a tumor suppressor-like candidate amenable to an overexpression or synthetic lethality strategy.

### Drug response correlation

To evaluate the therapeutic relevance of top-ranked inhibition targets, TF activity scores derived from CCLE breast cancer cell lines were correlated with drug IC50 values of the matching cell lines from the GDSC2 dataset. For inhibition targets, higher TF activity was expected to be associated with increased IC50 (reduced drug sensitivity), corresponding to a positive Spearman correlation direction consistent with BN predictions.

Among the top 10 inhibition targets, 8 showed directional consistency with BN predictions based on positive median Spearman correlation. Cell lines with higher target activity show higher IC50 values (lower sensitivity) across a broad range of drugs, suggesting that these targets confer widespread drug resistance and that its inhibition may restore sensitivity of multiple therapeutic agents. DNA damage response agents such as MK-1775 showed particularly strong associations with FOXA1 activity (ρ = +0.702, FDR = 3.59 × 10⁻⁶) (Figure 4c), consistent with its established role in mediating resistance to genotoxic stress in luminal breast cancer^36^. An analogous analysis for overexpression targets is shown with ETS1 with AZD7762 as a representative example (Figure 4d).

### Literature mining

The number of PubMed publications retrieved per TF across three categories of functional evidence, clinical relevance, and drug actionability was used to check the degree of prior characterization of each candidate (Figure 4e-f). Targets with high publication counts across all three categories were considered previously characterized, supporting the pipeline’s ability to recapitulate known biology. AR (999 drug, 1,000 clinical, 420 function) targeting drug, enzalutamide, has been reported to be effective on breast cancer^37^, and ongoing clinical trials are also reported^38^ . FOXA1 also showed high publication counts across all three categories (216 drug, 395 clinical, 113 function), consistent with its established role as a key regulator of estrogen receptor signaling. FOXA1 is known to be a pioneer factor that shapes chromatin accessibility for ESR1 and determines its downstream transcriptional program in luminal breast cancer^39^. Notably, FOXA1 ranked higher among the BN-predicted inhibition targets (rank 6) than ESR1 itself (rank 13, outside the top 10), and showed strong directional consistency with BN predictions in the GDSC2 drug response analysis, suggesting that the pipeline captures the broader regulatory influence of FOXA1 across cell line subtypes beyond the luminal-restricted activity of ESR1. Targets with low counts being flagged as potentially novel candidates with limited prior therapeutic investigation (Figure 4e–f).

## Discussion

In this study, we present BayesTx, a causal Bayesian network pipeline that integrates patient domain and cell line data to identify therapeutic transcription factors (TFs) in cancer. BayesTx utilizes TF activity scores (via DoRothEA or CollecTRI) and pathway scoring (via PROGENy or Hallmark). And by learning domain-specific causal graphs and merging them at an optimally weighted ratio, BayesTx produces a single consensus network. Using this network, the causal effect of individual TFs on cancer cell viability can be estimated through do-simulation. When applied to breast cancer, the pipeline identified a ranked list of TF targets and the predicted causal directions were supported by the external cohort validation using METABRIC survival data and GDSC2 drug response profiles for the majority of the suggested candidates.

This study has several limitations. First, the application of prior biological knowledge was limited. The blacklist constraints were done after the NOTEARS structure learning, rather than embedding them within the optimization process. And the whitelist constraints are designed to boost only near-zero edges after structure learning. This means that the learned graph structure does not strictly enforce prior knowledge. Also, gene level ACE extensions are derived indirectly through DoRothEA regulon propagation, and their interpretation as causal effects relies solely on the TF-target regulon relationships which limits the direct usage of the scores. Third, suggested targets were derived from pan-BRCA datasets, which does not consider the diverse nature of breast cancer subtypes. Future implementations could address these by using subtype-specific data as an input to build subtype-specific networks, whereas implementing blacklist and whitelist constraints directly into a continuous optimization process.

The BayesTx framework is designed to be applicable to a wide range of cancer and other diseases by substituting the domain-specific input data. The modular architecture supporting interchangeable pathway scoring methods (PROGENy or Hallmark) and TF activity methods (DoRothEA or CollecTRI) allows users to tailor the pipeline to the regulatory context of their specific disease of interest. Future studies include subtype-specific analysis, integration of additional omics layers such as metabolic network data, and direct incorporation of experimentally validated wet-lab results to close the loop between computational prediction and biological confirmation.

## Acknowledgements

This research was supported by the Ministry of Science and ICT, through the National Research Foundation of Korea (NRF) (RS-2023-00262527, RS-2025-25442169).

