## Supplementary 1 for "A Bayesian Network-Based Framework for Causal Cancer Drug Target Discovery Integrating Patient and Cell Line Data"

Supplementary 1. Blacklist and whitelist

Blacklist

| **Source** | **Target** |
|---|---|
| MAPK | EGFR |
| PI3K | EGFR |
| NFkB | JAK-STAT |

Whitelist

| **Source** | **Target** |
|---|---|
| EGFR | MAPK |
| EGFR | PI3K |
| TNFa | NFkB |
| VEGF | MAPK |
| VEGF | PI3K |
